# MSCquartets 1.0: Quartet methods for species trees and networks under the multispecies coalescent model in R

**DOI:** 10.1101/2020.05.01.073361

**Authors:** John A. Rhodes, Hector Baños, Jonathan D. Mitchell, Elizabeth S. Allman

## Abstract

`MSCquartets` is an R package for species tree hypothesis testing, inference of species trees, and inference of species networks under the Multispecies Coalescent model of incomplete lineage sorting. Input for these analyses are collections of metric or topological locus trees which are then summarized by the quartets displayed on them. Results of hypothesis tests at user-supplied levels are displayed in a simplex plot by color-coded points. The package includes the QDC and WQDC algorithms for topological and metric species tree inference, and the NANUQ algorithm for level-1 topological species network inference, all of which give statistically consistent estimators under the model.

## 1 Introduction

The advent of large multilocus datasets, often with discordant gene phylogenies, requires new statistical methods of species tree and network inference, with attention to attaining acceptable runtimes. The *Multispecies Coalescent* (MSC) model of incomplete lineage sorting (ILS), which is believed to be one of the most fundamental causes of gene tree discord, provides the theoretical framework for such work.

`MSCquartets` addresses the species tree/network inference problem through *quartet* summaries of gene tree collections. If *n* gene trees have been inferred on (subsets of) *N* taxa, the collection can be summarized by tabulating the counts of quartets displayed on the *n* trees for each of the 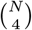 4-element subsets of the taxa. For example, if the four taxa {*a, b, c, d*} occur on *m* ≤ *n* of the gene trees, and the trees are fully resolved, then the counts *m*_*ab*|*cd*_, *m*_*ac*|*bd*_, *m*_*ad*|*bc*_ of the 3 possible displayed topologies can be tabulated, where *xy*|*zw* denotes the binary quartet tree separating *xy* from *zw*. The resultant *quartet count concordance factor qcCF* = (*m*_*ab*|*cd*_, *m*_*ac*|*bd*_, *m*_*ad*|*bc*_) or its normalization 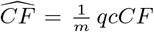, the *empirical quartet concordance factor* (Allman *et al*., 2020), are the data summaries of focus. The vector 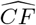 estimates the expectation *eCF* under the MSC of quartet frequencies displayed on gene trees. As the number *m* of gene trees sampled from the MSC grows, 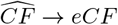 in probability.

While basic `MSCquartets` functions compute concordance factor summaries of multilocus datasets, the main analysis tools perform:

1. Hypothesis testing of *qcCF*s for fit of the MSC on a 4-taxon tree,
2. Inference of a large species tree from *qcCFs*,
3. Inference of a large topological species network from *qcCFs*.

Built on the `ape` and `phangorn` packages (Paradis and Schliep, 2018; Schliep, 2011), `MSCquartets` functions are designed for flexibility and easy export of output from R. Specialized graphics functions allow for display of hypothesis test results.

## 2 Methods

Data analysis with `MSCquartets` generally begins by tabulating qcCFs for a multilocus tree collection. This step’s running time of 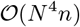 dominates that of the more substantive analyses 1-3 above, both theoretically and practically, but need only be performed once.

### 2.1 Hypothesis testing

Using new methodology developed by Mitchell *et al*. (2019), with applications discussed further by Allman *et al*. (2020), a *qcCF* can be tested for fit to the MSC on 4-taxon unrooted topological species trees.

Forms of the tests implemented in `MSCquartets` allow the null hypothesis *H*_0_: *The qcCF arose from a species quartet tree of specified topology under the MSC*, or *H*_0_: *The qcCF arose from a species quartet tree of unknown topology under the MSC*. With exhaustive alternative hypotheses, rejection of the null indicates in the first case that the MSC on the specific species tree topology is rejected, and in the second that the MSC on any tree is rejected (perhaps due to hybridization). A third test, using more routine statistical methods, can test the MSC on a quartet star tree (a polytomy) against any alternative, as has also been implemented by Sayyari and Mirarab (2018).

The primary hypothesis tests use a novel approximating distribution, dependent on the sample size *m*, that performs better than the standard asymptotic distribution for models such as these with singularities or boundaries. A simplex plot, such as in Figure 1 for a *Heliconius* dataset (Martin *et al*., 2013), displays test results for all 4-taxon subsets. For each of these, a color-coded symbol is plotted, with the black lines indicating all possible *eCFs* for the three 4-taxon species tree topologies. Blue circles, for example, indicate that a particular *qcCF* is consistent at the specified level with the MSC on a species tree, and lie in closer proximity to the model lines than the red triangles indicating rejection of the null hypothesis.

**Figure 1:**
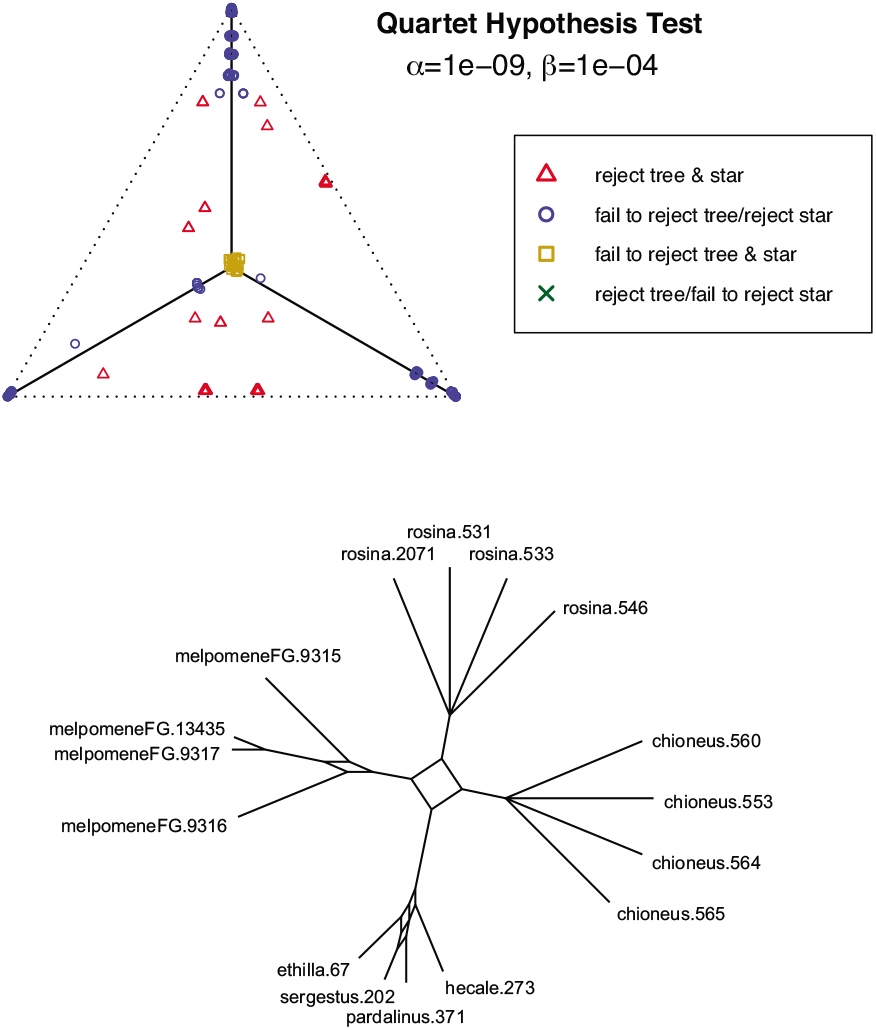
(Top) A simplex plot for *Heliconius* data shows results of two hypothesis tests. Rejection levels are *α* = 10^−9^ for a test of any tree topology under the MSC, and *β* = 10^−4^ for the polytomy test. With multiple individuals sampled for some taxa, failure to reject some polytomies is not surprising. (Bottom) The splits network constructed from the NANUQ distance for these data, presenting strong evidence for a 4-cycle in the species network.

### 2.2 QDC and WQDC species tree estimators

Unrooted species trees, either topological or metric, can be inferred from topological quartets displayed on gene trees by first computing novel intertaxon distances introduced by Rhodes (2019) and Yourdkhani and Rhodes (2020). `MSCquartets` includes functions to find these distances from tabulated *qcCFs*, allowing for rapid statistically-consistent inference under the MSC model, using standard tree building methods such as Neighbor Joining, which are already available in ape.

### 2.3 NANUQ species network topology estimator

The NANUQ algorithm for estimating a species network topology under the MSC was introduced by Allman *et al*. (2019). This fast method takes as input a collection of gene trees summarized by their 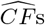, uses the outcome of the hypothesis tests discussed above to determine putative 4-taxon networks, and a modification to level-1 networks of the intertaxon quartet distance used by QDC to create a distance table. From the distance table, a splits graph is constructed after projecting to a circular split system with NeighborNet (Bryant and Moulton, 2004). Such a splits graph is shown for the *Heliconius* data in Figure 1. A last step involves interpretation of the splits graph using theoretically-derived rules to obtain an inferred topological species network, omitting any 2- and 3-cycles. In `MSCquartets`, a single function call accomplishes all steps up to the use of NeighborNet, which can be performed with `phangorn`, or externally to R with `SplitsTree` (recommended) (Huson and Bryant, 2006).

## 3 Conclusion

The R package `MSCquartets` implements a suite of fast and statistically consistent algorithms for species tree and network inference under the MSC model. In addition, it implements novel hypothesis testing methods for the fit of multilocus datasets to the MSC model. All of these are based on recent theoretical advances in the relationship between the topology of a tree or network and expected quartet concordance factors, providing a common statistical framework for the package.

The package is available on the CRAN, at https://CRAN.R-project.org/package=MSCquartets.

## Funding

This work was supported by the National Institutes of Health [R01 GM117590], under the Joint DMS/NIGMS Initiative to Support Research at the Interface of the Biological Mathematical Sciences, and [2P20GM103395], an NIGMS Institutional Development Award (IDeA).

## Notes

https://CRAN.R-project.org/package=MSCquartets

### Competing Interest Statement

The authors have declared no competing interest.

## References

Allman, E., Baños, H., and Rhodes, J. (2019). NANUQ: A method for inferring species networks from gene trees under the coalescent model. Algorithms Mol. Biol., 14(24), 1–25.

Allman, E., Mitchell, J., and Rhodes, J. (2020). Gene tree discord, simplex plots, and statistical tests under the coalescent. bioArXiv https://doi.org/10.1101/2020.02.13.948083.

Bryant, D. and Moulton, V. (2004). Neighbor-Net: An agglomerative method for the construction of phylogenetic networks. Molecular Biology and Evolution, 21, 255–265.

Huson, D. and Bryant, D. (2006). Application of phylogenetic networks in evolutionary studies. Mol. Biol. Evol., 23(2), 254–267.

Martin, S., Dasmahapatra, K., Nadeau, N., Salazar, C., Walters, J., Simpson, F., Blaxter, M., Manica, A., Mallet, J., and Jiggins, C. (2013). Genome-wide evidence for speciation with gene flow in Heliconius butterflies. Genome Res, 23, 1817–1828.

Mitchell, J., Allman, E., and Rhodes, J. (2019). Hypothesis testing near singularities and boundaries. Electron. J. Statist., 13(1), 2150–2193.

Paradis, E. and Schliep, K. (2018). ape 5.0: an environment for modern phylogenetics and evolutionary analyses in R. Bioinformatics, 35, 526–528.

Rhodes, J. (2019). Topological metrizations of trees, and new quartet methods of tree inference. IEEE/ACM Trans. Comput. Biol. Bioinform., early access.

Sayyari, E. and Mirarab, S. (2018). Testing for polytomies in phylogenetic species trees using quartet frequencies. Genes, 9(3), E132.

Schliep, K. (2011). phangorn: phylogenetic analysis in R. Bioinformatics, 27(4), 592—593.

Yourdkhani, S. and Rhodes, J. (2020). Inferring metric trees from weighted quartets via an intertaxon distance. arXiv:2002.04564.

